# Allosteric Ligand-Aptamer Complexes Orchestrate Supramolecular or Transient Catalytic, Transcription and Fibrinogenesis Processes

**DOI:** 10.1101/2025.10.26.684577

**Authors:** Diva Froim, Hadar Amartely, Jiantong Dong, Eli Pikarsky, Itamar Willner

## Abstract

Allosteric regulation, the modulation of biological macromolecular function through binding of molecules at distant sites distinct from the active site, is a fundamental principle in biology that governs enzyme activity, signaling, and gene expression. In this work, we present allosteric ligand/aptamer complexes, coupled to biocatalytic reaction modules composed of enzymes, DNAzymes, or transcription machineries, regulating the catalytic and transient functions of these frameworks. This principle is exemplified by the assembly of ligand/aptamer subunits supramolecular complexes that allosterically stabilize the Mg²⁺-dependent DNAzyme, allowing its ribonucleobase cleavage activity, promoting the formation of transcription templates that yield RNA products, and modulating the assembly of thrombin aptamer subunits that inhibit thrombin-induced coagulation. Specifically, melamine (Mel)/aptamer subunits complexes allosterically stabilize the assembly of Mg²⁺-dependent DNAzyme strands for substrate cleavage, the formation of thrombin aptamer subunits that inhibit the conversion of fibrinogen to fibrin, and the stabilization of a transcription template encoding the Malachite Green (MG) RNA aptamer. Furthermore, coupling an enzyme that depletes the ligand/aptamer complex, which allosterically stabilizes the biocatalytic reaction module, demonstrates the dissipative and transient operation of the catalytic system. This concept is illustrated by the adenosine (Ade)/aptamer subunits supramolecular complex, which stabilizes thrombin aptamer subunits to inhibit thrombin-induced fibrinogenesis, and promotes the formation of an active transcription template for RNA synthesis. In the presence of adenosine deaminase (ADA), Ade is transformed into inosine, which lacks affinity for the aptamer subunits, thereby degrading the Ade/aptamer assemblies and depleting the allosteric complexes. The temporal disassembly of these allosteric stabilizing complexes leads to the transient inhibition of thrombin-induced coagulation or to the transient operation of a transcription machinery.

For Table of Contents Only:

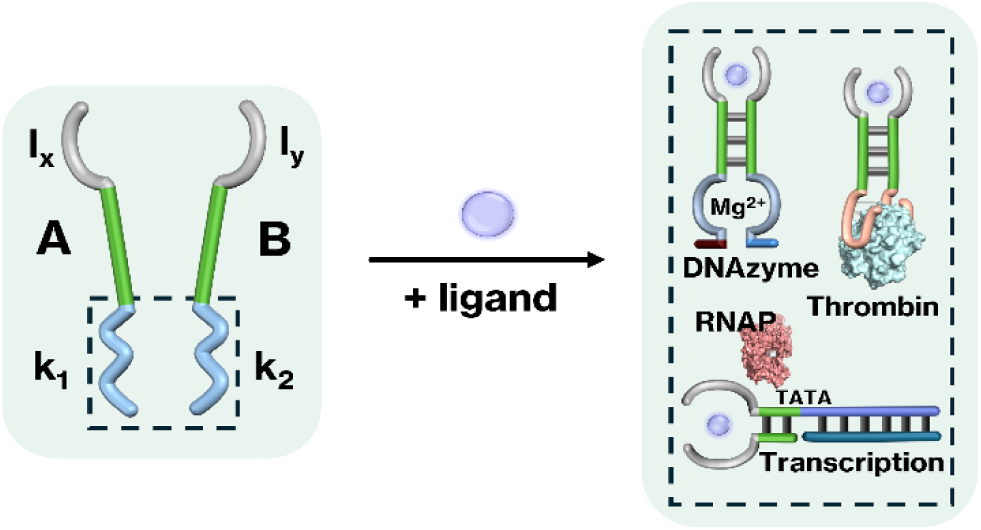

## Introduction

Allosteric regulation, the modulation of the activity of macromolecules through binding of molecules at distant sites, is a fundamental principle governing biological processes. The communication between spatially separated molecular domains, provides the dynamic flexibility that underlies metabolism, signaling, and gene expression.^1,2^ Jacques Monod famously referred to allostery as the “second secret of life,” emphasizing its central role in the self-organization and adaptability of biological systems.^3^ Introduction of the allostery concept into synthetic nucleic acid structures might add an important dimension to the functional properties of the biopolymer and expand its application in synthetic biology. The base sequence comprising nucleic acids encodes functional information into the structure of the biopolymer. Sequence guided functions of DNA include sequence-specific recognition and binding of biomolecules or low molecular-weight ligands (aptamers),^4-6^ sequence dictated displacement of duplex nucleic acids or protein/nucleic acid complexes by auxiliary strands,^7-9^ and sequence regulated catalytic properties in the presence or absence of auxiliary cofactors (DNAzymes or ribozymes), e.g., metal-ion or amino acid-dependent DNAzymes^10-13^ and hemin/G-quadruplex^14,15^ DNAzymes. Moreover, the sequences comprising DNA duplex frameworks dictate selective reactivity patterns towards auxiliary enzymes such as endonucleases^16,17^ or nickases.^18^ In addition, auxiliary enzymes such as DNA or RNA polymerases and added dNTPs or NTPs as fuels, catalyze in the presence of nucleic acid templates dictated polymerization and displacement of DNA or RNA products.^19,20^ This arsenal of recognition and catalytic functions embedded in oligonucleotides provides a versatile "tool-box" for the rapidly developing area of DNA nanotechnology.^21^ Over the years, the functional information embedded in nucleic acids has been implemented to develop DNA switches,^22,23^ machines^24,25^ and two- and three-dimensional DNA nanostructures.^26-28^ In addition, dynamically reconfigured nanostructures,^29-31^ programmed logic gate circuits,^30,31^ dynamic reconfigurable DNA networks,^32^ dissipative circuits,^33-35^ and switchable transcription machineries^36,37^ were demonstrated. Moreover, chemical modifications of aptamers with light responsive^38^ or redox active units^39^ led to switchable binding properties of aptamers. Conjugation of aptamers to DNAzyme catalytic units yielded hybrid structures, "nucleoaptazymes", emulating native enzymes by providing cooperative substrate binding sites in spatial proximity to the active site in the conjugated structure.^40,41^ Diverse applications of aptamers were demonstrated, including their use as sensing^42,43^ and imaging materials,^44^ engineering of stimuli-responsive drug-carriers and their targeting to specific cell receptors.^45,46^ Also, aptamers were used as therapeutic agents through selective binding to proteins and their inhibition, e.g., association to VEGF (Inhibiting angiogenesis)^47,48^ or thrombin (Inhibiting fibrinogenesis).^49^ Similarly, DNAzymes have found broad applications as amplifiers of sensing events, using *in vitro* or *in vivo* assays^50^ and catalyzing diverse chemical transformations, such as oxidation of NADH^51^ or dopamine.^52^ Also, DNAzymes were employed as synthetic catalysts for gene therapy^53^ and for the generation of reactive oxygen species for chemo dynamic cancer therapy.^54,55^

Here we wish to report on the conjugation of ligand/aptamer supramolecular complexes to DNAzyme subunits, proteins (thrombin) and transcription templates, resulting in the allosteric operation of a DNAzyme, inhibition of fibrinogenesis and RNA transcription. Melamine (Mel) or adenosine (Ade) act as ligands assembling ligand/aptamer supramolecular complexes that allosterically guide the respective catalytic, fibrinogenic and transcription circuits. By coupling adenosine deaminase (ADA) to the Ade aptamer allosterically-stabilized fibrinogenesis and transcription circuits, transient, dissipative, operation of the frameworks is achieved. Beyond the expansion of the functionalities of stimuli-responsive nucleic acid circuits, the significance of the ligand/aptamer complex, allosterically-driven, catalytic circuits is reflected by: (i) their possible application for sensing (e.g., Mel); (ii) the temporal, dose-controlled, therapeutic applications of the circuits (e.g., inhibition of thrombin-induced fibrinogenesis); and (iii) the spatiotemporal control-over transcription machineries by synthetic ligand/aptamer complexes emulating functions of native transcription factors.

## Results and Discussion

The general concept to allosterically operate catalytic circuits consisting of DNAzymes, proteins (thrombin) or transcription machineries by ligand/aptamer complexes is exemplified in Figure 1A using melamine (Mel)/aptamer complexes.^56,57^ The system consists of two strands A_m_ and B_m_ that include engineered sub-domains l_1_ and l_2_ corresponding to aptamer subunits of the ligand conjugated to the catalytic subunits k_1_ and k_2_. The catalytic subunits include the pre-engineered sequences to operate a DNAzyme, thrombin binding or transcription machinery catalytic functions. While the strands A_m_ and B_m_ have partial complementarity, it is insufficient to form a stable interstrand complex. However, binding of the ligand (Mel) to the aptamer subunits results in an interstrand supramolecular complex cooperatively stabilized by the ligand/aptamer complex and the complementarity associated with the two strands. Furthermore, the spatial proximity between the tethers k_1_ and k_2_ is pre-engineered to evolve the catalytic function in the supramolecular assembly. That is, the ligand-induced formation of ligand/aptamer supramolecular complex allosterically stabilizes and activates the catalytic function of the spatially confined k_1_ and k_2_ subunits.

**Figure 1.**
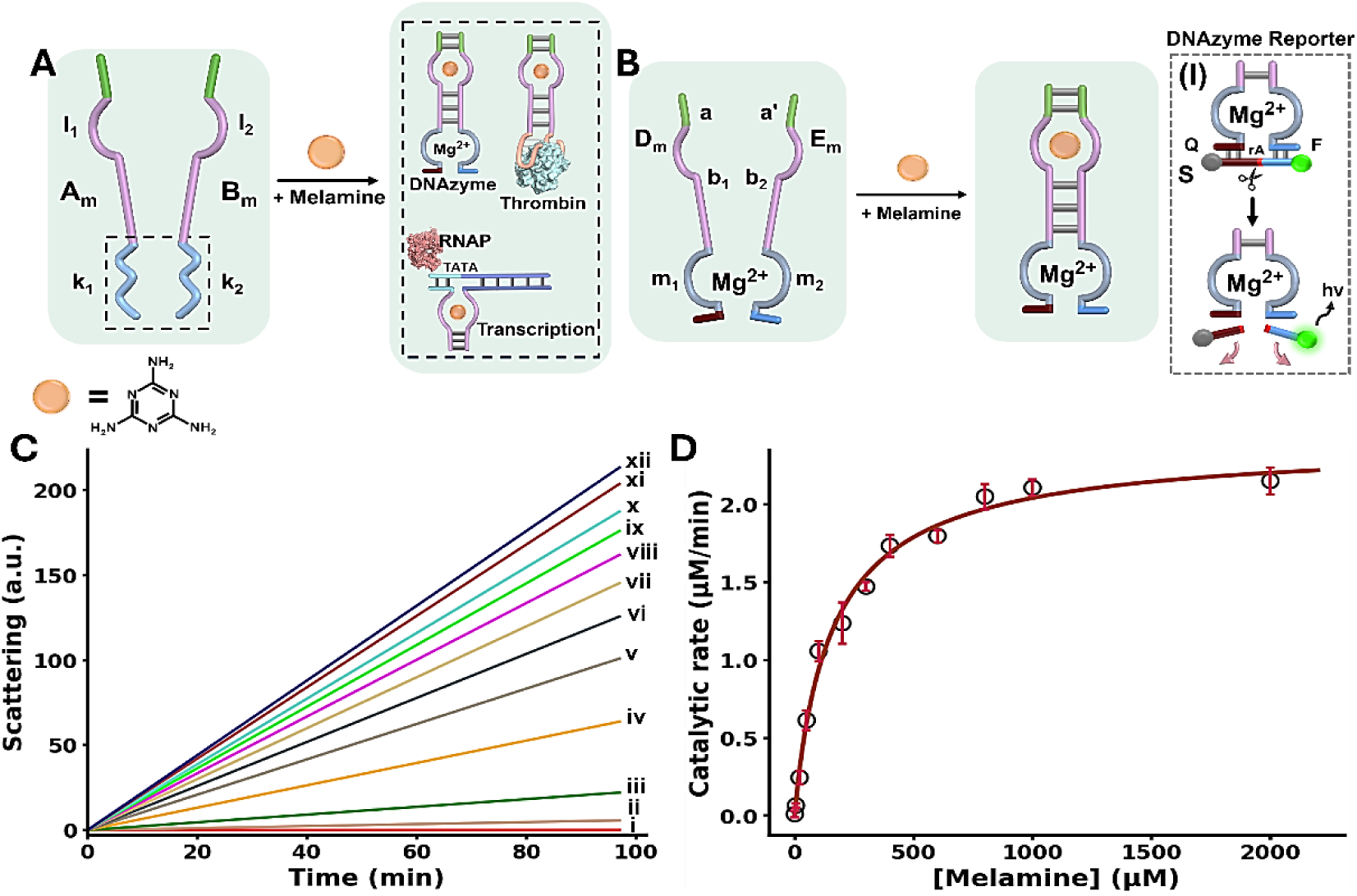
(A) Allosteric activation of catalytic DNA circuits by supramolecular assembly of ligand/aptamer subunits complexes using melamine (Mel)/aptamer subunits. (B) Schematic Mel-induced allosteric stabilization of a Mg^2+^-ion dependent DNAzyme through Mel/aptamer subunits complex. Panel I – Probing the catalytic activity of the DNAzyme by the DNAzyme-catalyzed ribonucleobase cleavage of a fluorophore (FAM)/quencher (BHQ1)-modified substrate strand, S. (C) Time-dependent fluorescence changes upon cleavage of the F/Q-modified substrate by the allosterically-stabilized DNAzyme formed in the presence of variable concentrations of Mel: (i) 0 µM, (ii) 5 µM, (iii) 20 µM, (iv) 50 µM, (v) 100 µM, (vi) 200 µM, (vii) 300 µM, (viii) 400 µM, (ix) 600 µM, (x) 800 µM, (xi) 1000 µM, (xii) 2000 µM. (D) Catalytic rates of the allosterically stabilized DNAzyme, in the presence of variable concentrations of Mel (Fluorescence intensities were translated to free fluorophore concentration using the appropriate calibration curve, Figure S1).

Figure 1B schematically depicts the Mel/aptamer subunits complex induced allosteric stabilization of a supramolecular complex activating the function of a Mg^2+^-ion dependent DNAzyme. The reaction circuit includes two strands D_m_ and E_m_ which include subunits b_1_ and b_2_ of the split Mel aptamer, and subunits m_1_ and m_2_ corresponding to the arm/loop sequences of the DNAzyme. The complementary sequence domains a and a’ were added to strands D_m_ and E_m_. In the presence of Mel, the supramolecular complex cooperatively stabilized by the Mel/aptamer subunits complex and the interbridging duplex a/a’ is formed, resulting in the spatially confined DNAzyme configuration composed of m_1_ and m_2_. The allosteric stabilization of the functional Mg^2+^-ion dependent DNAzyme is then probed by the cleavage of the fluorophore/quencher modified substrate S by the DNAzyme, panel I (F = FAM; Q = BHQ1).

Figure 1C, depicts the rate of cleavage of the F/Q-modified substrate by the allosterically, Mel-stabilized DNAzyme framework in the presence of variable Mel concentrations. While no cleavage of the substrate proceeds in the absence of Mel, consistent with the lack of communication between strands D_m_ and E_m_, curve i, the supramolecular DNAzyme structure is activated in the presence of Mel, curves ii-xii.

As the concentration of Mel increases, the rate of cleavage is enhanced. Figure 1D, displays the rates of cleavage of the substrate strand in the presence of variable Mel concentrations. A saturation curve is observed consistent with the saturated formation of the Mel/aptamer subunits complex from which the V_max_ = 2.41 ±0.07 µmol⋅min^-1^ and K_0.5_ = 162 ±4.8 µM for the allosterically-stabilized supramolecular DNAzyme were evaluated. The detection limit of the DNAzyme was calculated to be 9.3 ±0.3 nM using the three-sigma method. Isothermal titration calorimetry (ITC) experiments supported the Mel/aptamer subunits formation of the supramolecular DNAzyme structure revealing a K_d_ of 0.890 ±0.123 µM corresponding to the complex between Mel and the two subunits D_m_ and E_m_, Figure S2. Beyond demonstrating the Mel/aptamer induced allosteric activation of the Mg^2+^-ion dependent DNAzyme, the system presents as an amplified Mel sensing platform. Mel sensing is of importance as Mel has been used as an illegal ingredient additive in food products causing severe health problems in infants.^58^ Different analytical methods including mass-spectrometry,^59^ high performance liquid chromatography^60^ and CRISPR/Cas14a sensing platforms^61^ were developed to detect Mel residues in food products. The amplified Mel-dependent operation of the DNAzyme allows the analysis of Mel with a detection limit corresponding to 0.001 ppm, that is lower than the detection threshold defined by the FDA (1.0 ppm).^62,63^

The allosteric Mel/aptamer subunits complex stimulating the catalytic functions of a conjugated DNA framework were further demonstrated with the Mel-mediated inhibition of fibrinogen to fibrin coagulation.

Thrombin is a key physiological regulator of the blood clotting mechanism.^64^ While it plays a key role in hemostatic clotting of vascular injuries, its balanced dose activity is crucial to prevent blood clots and thrombosis.^65,66^ Diverse anti-coagulant therapeutic agents, controlling thrombin activity are known.^67,68^ Within these efforts, anti-thrombin aptamers that bind to thrombin were isolated and their inhibition of thrombin was implemented to design anti-coagulant agents.^69,70^ The dose-controlled, inhibition of thrombin catalytic functions by aptamers, acting as an anti-coagulation agents, could thus be a significant advance in controlling blood clotting (thrombosis). Accordingly, the allosteric activation of the blood clotting capacities of the anti-thrombin aptamer framework using an auxiliary ligand, e.g. Mel, could be an interesting path to follow. This concept is exemplified in Figure 2A with the design of a Mel/aptamer subunits allosteric circuit for the controlled inhibition of thrombin’s coagulation function.

**Figure 2.**
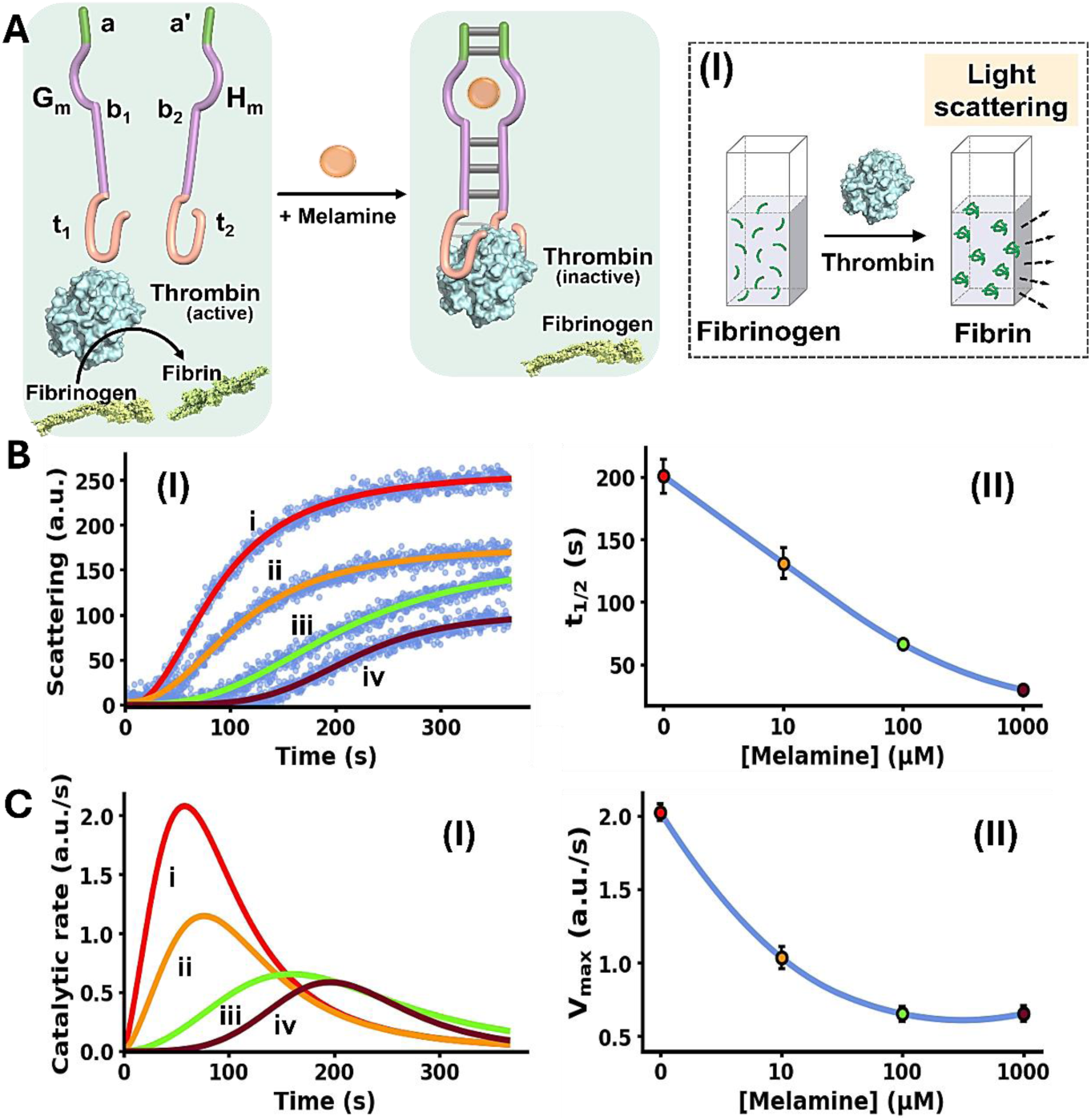
(A) Schematic melamine (Mel)-induced allosteric inhibition of thrombin-induced fibrinogenesis of fibrinogen to fibrin through the formation of thrombin/thrombin aptamer subunits framework. Panel I – Schematic probing of the fibrinogenesis process by dynamic light-scattering. (B) Panel I – Temporal light-scattering curves upon inhibition of thrombin-induced fibrinogenesis of fibrinogen to fibrin in the presence of the thrombin/Mel aptamer subunits framework G_m_/H_m_ and variable concentrations of Mel: (i) 0 µM, (ii) 10 µM, (iii) 100 µM, (iv) 1000 µM. Panel II – t_1/2_ values derived from fibrinogenesis temporal light-scattering curves, in the presence of variable concentrations of Mel. (C) Panel I – Temporal catalytic rates of thrombin-induced fibrinogenesis in the presence of G_m_/H_m_ strands, allosterically stabilized by variable concentrations of Mel: (i) 0 µM, (ii) 10 µM, (iii) 100 µM, (iv) 1000 µM. Panel II – Maximum fibrinogenesis rates (V_max_) upon subjecting G_m_/H_m_ to variable concentrations of Mel. In all experiments, G_m_/H_m_ = 1.0 µM, thrombin = 5 nM, fibrinogen = 10 mg/ml. Data are means±SD, N=3.

The system consists of two strands G_m_ and H_m_ composed of the subsequences b_1_ and b_2_ corresponding to the Mel aptamer subunits, extended by four base complementary tethers, a and a’, cooperatively enhancing the stability of the Mel/aptamer subunits complex. The Mel aptamer subunits G_m_ and H_m_ are further extended by the tethers t_1_ and t_2_ that correspond to the anti-thrombin aptamer subunits. Despite the partial complementarity of strands G_m_ and H_m_ and the binding affinity of thrombin to the subunits t_1_ and t_2_, the strands G_m_ and H_m_ lack binding affinity to allow the formation of the G_m_/H_m_/thrombin complex, and thus, the inhibition of thrombin-induced fibrinogenesis (coagulation of fibrinogen to fibrin) is prohibited. The rate of thrombin-induced coagulation in the absence of the strands G_m_/H_m_ yet in the presence of Mel, is reflected by the temporal light-scattering of the system, show similar temporal light-scattering intensity changes to free thrombin in the absence of ligands, Figure S3. The allosteric Mel/aptamer subunits stabilization of the thrombin/anti-thrombin subunits results in the Mel-induced, controlled inhibition of the thrombin-induced coagulation of fibrinogen to fibrin. The rate of fibrinogenesis is followed by the temporal light-scattering features associated with the coagulation of fibrinogen to fibrin, in the presence of variable Mel concentrations, Figure 2A, panel I. Figure 2B, panel I, depicts the time-dependent light-scattering intensity changes associated with the thrombin-induced coagulation of fibrinogen to fibrin in the presence of strands G_m_ and H_m_, in the absence of Mel, curve (i), and in the presence of variable concentrations of Mel, curves (ii)-(iv). While in the absence of Mel rapid fibrinogenesis is observed, the addition of Mel induces the formation of interstrand Mel/G_m_/H_m_ and the coagulation of fibrinogen to fibrin is suppressed, thus as the concentration of Mel increases the degree of inhibition of the fibrinogenesis is higher. The temporal light-scattering curves probing the allosteric inhibition efficacy of fibrinogenesis, in the presence of variable concentrations of Mel were quantitatively evaluated using two parameters.^71,72^ One parameter, t_1/2_, is the time interval corresponding to the light-scattering intensity reaching 50% of the saturation value at variable concentrations of Mel. Figure 2B, panel II, depicts the relation of t_1/2_ to the concentrations of Mel inducing allosterically the inhibition of thrombin. As the concentration of Mel increases the t_1/2_ light-scattering intensity values are lower reflecting an enhanced thrombin inhibition capability of the circuit. A second parameter evaluating the Mel-induced inhibition of thrombin is the maximum coagulation rates of fibrinogen to fibrin (V_max_) that are derived from the temporal light-scattering curves in the presence of variable Mel concentrations, first order time dependent derivative curves are shown in Figure 2C, panel I. The V_max_ values of the circuit, characterizing the inhibition efficiency induced by Mel, derived from the temporal light-scattering curves shown in Figure 2B, panel I, are displayed in Figure 2C, panel II. While a high V_max_ in the absence of Mel is observed, reflecting low thrombin inhibition, the V_max_ values decrease as the concentration of Mel increases, demonstrating the enhanced efficiency of Mel-induced inhibition of coagulation of fibrinogen to fibrin. The results displayed in Figure 2 introduce a new paradigm for controlling thrombin-induced coagulation by employing an auxiliary ligand (Mel) that allosterically regulates the dose-controlled formation of the anti-thrombin aptamer subunits/thrombin affinity complex that inhibits the coagulation process.

In the next step, the allosteric Mel-induced activation of a transcription machinery was examined. Regulation of RNA transcription controls many biological processes ranging from cell cycle progression^73^ and maintenance of intracellular metabolism to cellular differentiation.^74^ The transcription apparatus demonstrates dynamic adaptive features, primarily modulated by transcription factors.^75-78^ Beyond the key functions of the native transcription machinery in maintenance of living organisms, misregulation of transcription programs by dysfunctional transcription factors is the origin of various diseases including cancer, viral infection, neurological disorders, autoimmune pathologies and diabetes.^79,80^ Development of biomimetic synthetic transcription circuits is important not only to emulate the native apparatus by artificial model systems, but it could provide versatile therapeutic applications. For example, the controlled programmed *in vivo* synthesis of pre-engineered RNA could be a valuable source of therapeutic agents, e.g. aptamers, siRNAs and ribozymes. Indeed, recent research efforts demonstrated the modulation of transcription machineries by topological nucleic acid barriers conjugated to transcription templates, such as G-quadruplexes or DNA triplexes modeling native transcription factors’ functions.^71^ In the forthcoming section we introduce the allosteric Mel-induced operation of a transcription circuit as a biomimetic model system emulating the functions of transcription factors.

The Mel/aptamer subunits complex triggered activation of the transcription machinery is schematically displayed in Figure 3A. The inactive reaction module consists of the template strands N_m_/T_m_ containing an incomplete T7 RNAP promoter, the strand P_m_ and T7 RNA polymerase/NTPs mixture. The strands T_m_ and P_m_ include tethers b_1_ and b_2_ corresponding to the Mel aptamer subunits, where b_1_ is extended by the sequence x’ that is complementary to domain x in the template N_m_/T_m_. While x’ contains the sequence to complete the promoter region that activates the N_m_/T_m_ transcription machinery, the stability of the complementary duplex x/x’ is, however, insufficient to activate the transcription machinery. In the presence of Mel, the cooperative formation of the Mel/aptamer subunits complex, and the duplex x/x’ form an energetically stabilized, promoter-activated, transcription template enabling the activation of the transcription machinery, resulting in the T7 catalyzed RNAP/NTPs transcription of the RNA product, R_1_. The template N_m_/T_m_ is pre-engineered to yield the Malachite Green (MG) RNA aptamer as the transcription product. The resulting fluorescent MG/RNA aptamer complex (λ_ex_= 632nm; λ_em_= 650nm) provides, then, an optical readout signal for the temporal operation of the transcription machinery. Figure 3B depicts the time dependent fluorescence change caused by the production of the MG aptamer RNA product, generated in the absence of Mel, curve i, and in the presence of variable concentrations of Mel, curves ii-vi. Using an appropriate calibration curve, relating the fluorescence intensity of MG/RNA aptamer to its concentration, Figure S4, the rates of MG/RNA aptamer formation as a function of Mel concentrations were evaluated as depicted in Figure 3C. Peak rates (V_max_) of the transcription template derived from Figure 3C are displayed in Figure 3D, as Mel concentration increases, V_max_ increases.

**Figure 3.**
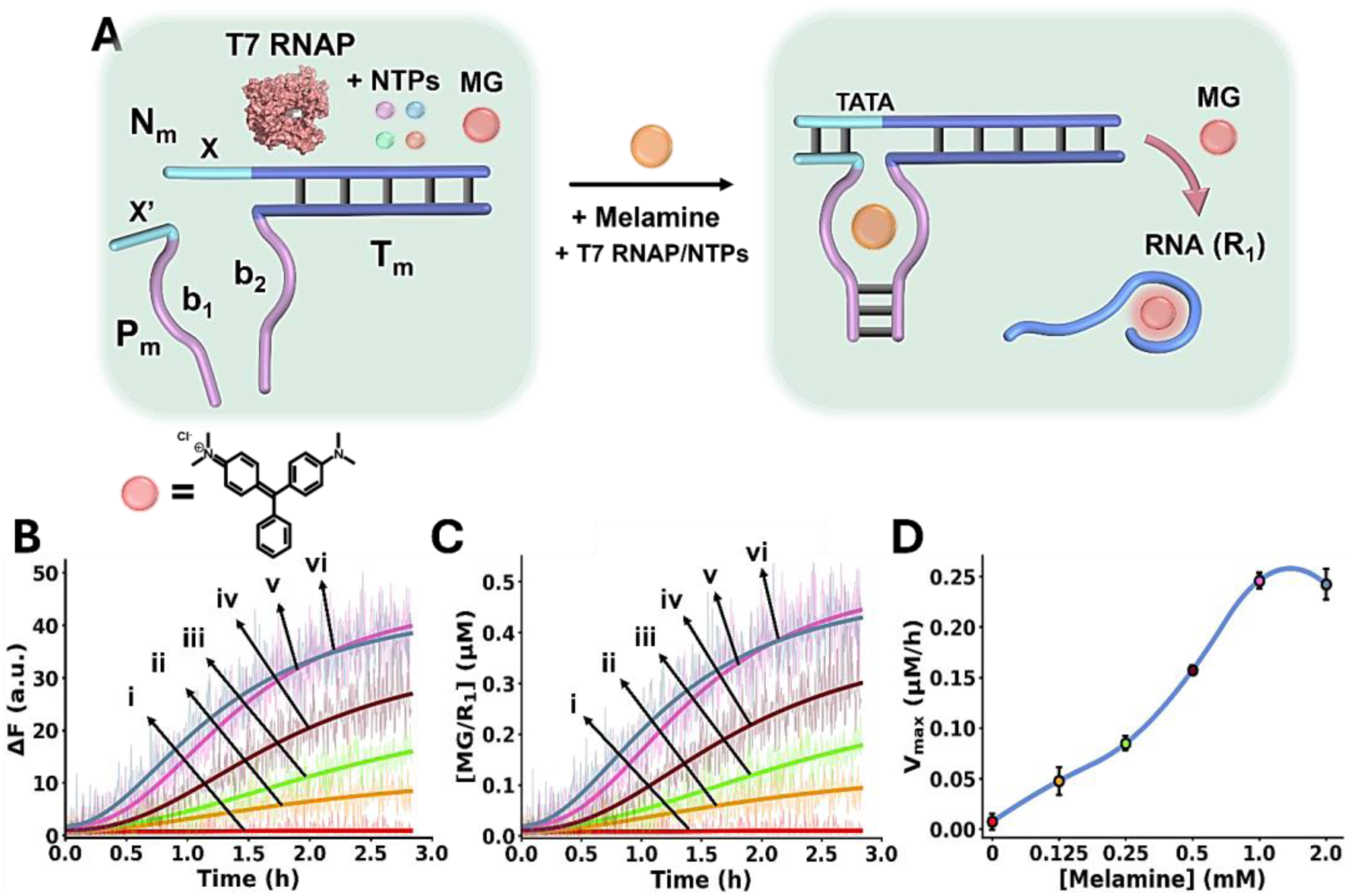
(A) Schematic melamine (Mel)/aptamer subunits allosterically triggering of the transcription machinery transcribing the Malachite Green (MG) RNA aptamer. The fluorescent MG/aptamer complex provides the readout signal for the transcription process. (B) Time-dependent fluorescence changes of the MG/RNA aptamer transcribed product generated in the presence of variable concentrations of Mel: (i) 0 mM, (ii) 0.125 mM, (iii) 0.25 mM, (iv) 0.5 mM, (v) 1.0 mM, (vi) 2.0 mM. (C) Temporal concentration changes of the MG/RNA aptamer transcribed product generated in the presence of variable concentrations of Mel: (i) 0 mM, (ii) 0.125 mM, (iii) 0.25 mM, (iv) 0.5 mM, (v) 1.0 mM, (vi) 2.0 mM (Translation of the temporal fluorescence changes shown in (B) to MG/aptamer concentrations were performed using the calibration curve provided in Fig. S4). (D) Maximum catalytic rates (V_max_) corresponding to the formation of the transcribed MG/aptamer product, in the presence of variable Mel concentrations. In all experiments, N_m_/T_m_ = 0.2 µM, P_m_ = 0.2 µM, NTPs = 0.5 mM, T7 RNAP = 1.5 U·µl^-1^. Data are means±SD, N=3.

The systems discussed so far demonstrated the allosteric ligand (Mel)-induced activation of catalytic processes involving a DNAzyme, thrombin-induced fibrinogenesis or a transcription machinery. Many of the catalytic processes in nature are, however, temporally modulated, leading to dissipative, transient and out-of-equilibrium operation.^81^ The ligand/aptamer complex allosterically regulating catalytic processes introduced a mechanism controlling the "dose" of the catalytic transformation. The coupling of a mechanism modulating temporally and transiently the allosteric mechanism could introduce an additional dimension to the "dose" regulated control over catalytic processes. The principle of dissipative out-of-equilibrium systems involves the design of reaction circuits that are activated by an auxiliary energy-fueled input (chemical fuel, light, electrical or magnetic stimuli) that generate an intermediate out-of-equilibrium state. The system includes, however, an internal mechanism depleting the auxiliary energy-fueled input resulting in the degradation of the intermediate state, into waste products while recovering the parent circuit. This leads to the temporal, transient, formation and depletion of the intermediate state. Substantial recent research efforts addressed the use of nucleic acid-based frameworks as functional reaction modules to design transient DNA circuits.^82,83^ Different triggers including nucleic acid fuel strands or light coupled to enzymes or DNAzymes were employed to trigger the temporal transitions of DNA frameworks into intermediate states that are temporally depleted by the catalysts to the parent reaction modules, thereby establishing transient operating DNA circuits.^84-86^ Diverse applications of transient operating circuits were demonstrated, including transient operating biocatalytic cascades,^87,88^ transient DNA-based load-release systems,^89^ or transient nucleic acid guided aggregation/de-aggregation of metal nanoparticles or semiconductor quantum dots.^90^ The allosteric ligand/aptamer stabilization of catalytic frameworks, and the availability of enzymes degrading the ligands, suggests that coupling of enzymes to the allosterically ligand/aptamer stabilized catalytic frameworks could provide a versatile mechanism to engineer dynamic, transient allosterically ligand-stabilized reaction modules. For example, diverse ligand/aptamer complexes can be coupled with ligand degrading enzymes (e.g., adenosine/adenosine deaminase, acetylcholine/acetylcholinesterase, uric acid/uricase) resulting in the separation of the complexes, as schematically shown in Figure S5.

The conjugation of an enzyme to an allosteric ligand-induced catalytic transformation leading to transient allosterically-driven catalytic process is schematically presented in Figure 4A, using the Ade/aptamer subunits complex and adenosine deaminase (ADA) as regulators controlling allosteric catalytic processes such as fibrinogenesis or transcription. The two strands A_a_ and B_a_ coupled with ADA, act as the reaction module. The strands A_a_ and B_a_ include the Ade aptamer subunits l_3_ and l_4_ conjugated to strands k_1_ and k_2_ that encode the nucleic acid sequences comprising the catalytic frameworks. Fueling the system with Ade results in the formation of Ade/aptamer subunits supramolecular complex allosterically stabilizing the catalytic framework consisting of the anti-thrombin aptamer subunits/thrombin fibrinogenesis inhibiting complex, or the active transcription machinery as intermediate products. The ADA present in the system concomitantly transforms Ade to inosine, that lacks affinity towards the aptamer subunits. Separation of the Ade/Ade aptamer complex recovers the parent reaction circuit, in which the catalytic transformations are prohibited. This leads to the transient Ade/ADA allosteric operation of fibrinogenesis or transcription processes. Indeed, in a recent study, the Ade/aptamer subunits stabilized Mg^2+^-ion DNAzyme was integrated with ADA in liposome protocells,^91^ and its transient operation in the cell-like reservoir was demonstrated. In the forthcoming section, the transient Ade/ADA operation of transcription and fibrinogenesis machineries will be addressed.

**Figure 4.**
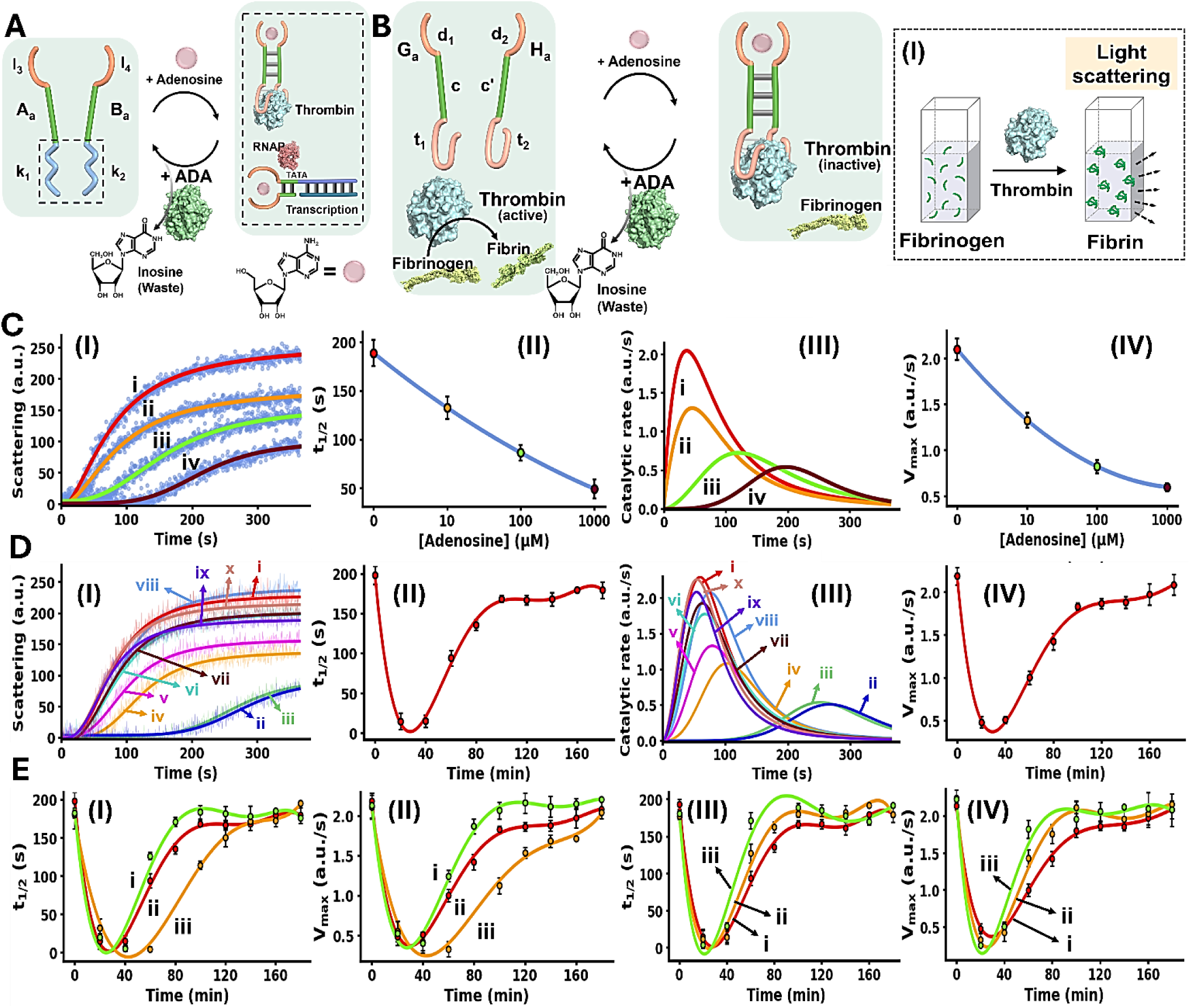
(A) Allosteric transient activation of catalytic DNA systems by supramolecular assembly of ligand/aptamer subunits complexes using adenosine (Ade)/Ade aptamer and adenosine deaminase (ADA). (B) Schematic application of Ade/aptamer subunits for the Ade/ADA transient allosteric inhibition of thrombin-induced fibrinogenesis. Panel I – Probing thrombin activity by following temporal light-scattering intensities associated with coagulation of fibrinogen to fibrin. (C) Panel I – Light-scattering intensities upon coagulation of fibrinogen to fibrin using the reaction circuit shown in (B) operating in the presence of variable Ade concentrations, yet in the absence of ADA: (i) 0 µM, (ii) 10 µM, (iii) 100 µM, (iv) 1000 µM. Panel II – t_1/2_ values of the reaction circuit in the presence of variable Ade concentrations extracted from panel I. Panel III – Evaluation of the catalytic rates associated with the temporal light-scattering intensity changes, in the presence of variable concentrations of Ade: (i) 0 µM, (ii) 10 µM, (iii) 100 µM, (iv) 1000 µM. Panel IV – Maximum catalytic rates (V_max_) associated with the system’s operation shown in panel III. (D) Panel I – Temporal light-scattering intensities corresponding to samples withdrawn at time-intervals, from the reaction circuit displayed in (B) demonstrating transient fibrinogenesis inhibition capability in the presence of Ade = 1.5 mM; ADA = 0.045 U·ml^-1^, after: (i) 0, in the absence of Ade, (ii) 20, (iii) 40, (iv) 60, (v) 80, (vi) 100, (vii) 120, (viii) 140, (ix) 160, (x) 180 minutes. Panel II – Analysis of temporal t_1/2_ values corresponding to transient fibrinogenesis induced by the reaction module shown in (B), Ade = 1.5 mM; ADA = 0.045 U·ml^-1^. Panel III – Analysis of the temporal catalytic rates of the temporal light-scattering intensities shown in panel I, of the transient fibrinogenesis induced by the reaction circuit displayed in (B), Ade = 1.5 mM; ADA = 0.045 U·ml^-1^. Panel IV – Transient V_max_ values derived from panel III. (E) Panels I and II – Transient fibrinogenesis driven in the presence of ADA = 0.045 U·ml^-1^ and different concentrations of Ade: (i) 1.25 mM, (ii) 1.5 mM, (iii) 1.8 mM, displayed using t_1/2_ and V_max_ parameters. Panels III and IV – Transient fibrinogenesis driven in the presence of Ade = 1.5 mM and different concentrations of ADA: (i) 0.045 U·ml^-1^, (ii) 0.055 U·ml^-1^, (iii) 0.065 U·ml^-1^, displayed using t_1/2_ or V_max_ parameters. In all experiments, G_a_/H_a_ = 1.0 µM, thrombin = 5 nM, fibrinogen = 10 mg/ml. Data are means±SD, N=3.

Figure 4B depicts schematically the reaction circuit used for the allosteric transient inhibition of the thrombin-induced coagulation of fibrinogen to fibrin in the presence of Ade and ADA. The system consists of two strands G_a_ and H_a_ as the DNA functional framework and ADA as the auxiliary catalyst. The strands G_a_ and H_a_ include the Ade aptamer subunits d_1_ and d_2_ conjugated to the complementary sequences c and c’ and extended by the anti-thrombin aptamer subunits t_1_ and t_2_. While in the absence of Ade, the base complementarity of c and c’ is insufficient to form a stable G_a_/H_a_ duplex structure that binds to thrombin and inhibits the thrombin-induced fibrinogenesis. The addition of Ade leads to the formation of the Ade/aptamer subunits complex, cooperatively stabilized by the c/c’ duplex, leading to the assembly of the supramolecular G_a_/H_a_ complex that stabilizes the thrombin aptamer subunits allowing the association of the interstrand Ade/G_a_/H_a_ supramolecular thrombin aptamer subunits framework to thrombin. Binding of the interstrand Ade/G_a_/H_a_ to thrombin allosterically inhibits thrombin-induced fibrinogenesis. The ADA coupled to the reaction circuit, concomitantly transforms Ade to inosine, leading to the separation of the G_a_/H_a_ units from thrombin, thereby recovering the free thrombin, exhibiting non-inhibited coagulation rate. That is, the system reveals transient allosteric Ade-induced inhibition of thrombin-induced coagulation of fibrinogen to fibrin. The degree of inhibition is controlled by the concentration of Ade that regulates the allosteric formation of the Ade/aptamer subunits complex. While the ADA concentration dictates the rate of recovery of the parent module, showing non-inhibited fibrinogenesis. That is, the transient allosteric inhibition of thrombin is regulated by two parameters; the concentration of Ade that triggers the fibrinogenesis inhibition and the concentration of ADA degrading Ade thereby regulating the temporal depletion of the inhibition phenomenon. The temporal and transient allosteric inhibition of thrombin, is then, followed by the temporal light-scattering features associated with the coagulation of fibrinogen to fibrin, in the presence of variable Ade and ADA concentrations, Figure 4B, panel I. The rate of thrombin-induced coagulation in the absence of the strands G_a_/H_a_, yet in the presence of Ade, show similar temporal light-scattering intensity changes to free thrombin in the absence of ligands, Figure S3.

Figure 4C, panel I, shows the temporal light-scattering intensities associated with the fibrinogenesis in the presence of different concentrations of Ade, yet in the absence of ADA: (i) 0 µM (ii) 10 µM (iii) 100 µM (iv) 1000 µM. As the concentration of Ade increases, the allosteric inhibition of the thrombin-induced fibrinogenesis increases as reflected by a prolonged initial lag and lower saturation values of temporal light-scattering curves. The analysis of the temporal light-scattering curves in panel I, in terms of t_1/2_, temporal catalytic rates and V_max_ values are presented in Figure 4C panels II-IV. Figure 4D, panel I shows the temporal light-scattering curves associated with the thrombin-induced fibrinogenesis at time-intervals of the system’s operation using Ade = 1.5 mM; ADA = 0.045 U·ml^-1^. While the control of the system, curve (i), in the absence of Ade demonstrates non-inhibited fibrinogenesis, addition of Ade leads to effective inhibition of fibrinogenesis, curve (ii), The system reveals, however, a continuous temporal change in the efficiency of fibrinogenesis reflected by the decrease in the inhibition effect in the system and ultimately shows the recovery of the non-inhibited fibrinogenesis behavior of the system, curves (ii)-(x). Analysis of the temporal light-scattering curves shown in Figure 4D panel I in terms of t_1/2_, temporal catalytic rates and V_max_ are summarized in Figure 4D, panels II-IV. The result demonstrates the transient allosteric Ade-induced inhibition of thrombin by Ade/ADA in the G_a_/H_a_ reaction circuit. Figure 4E depicts the effects of different Ade and ADA concentrations on the transient allosteric inhibition of thrombin-induced fibrinogenesis, as reflected by the t_1/2_ and V_max_ values derived from the temporal light-scattering intensities in the respective G_a_/H_a_ operating system. As the concentration of Ade increases, at fixed ADA concentration 0.045 U·ml^-1^, the time-interval of the transient recovery of the allosterically inhibited thrombin-induced fibrinogenesis is prolonged, Figure 4E, panels I and II. In addition, as the ADA concentration increases, at a fixed Ade concentration 1.5 mM, the time-interval of the transient recovery of the non-inhibited thrombin-induced fibrinogenesis is shortened, Figure 4E, panels III and IV. These results are consistent with the allosteric transient inhibition of the thrombin catalyzed coagulation of fibrinogen to fibrin regulated by the Ade/ADA-G_a_/H_a_ reaction circuit.

An additional process demonstrating the transient allosteric activation of a biocatalytic process included the Ade/ADA allosteric transient activation of a transcription machinery. Beyond transcription factor-modulated transcription machineries and accompanying regulated gene expression, promoter control elements regulate the dynamic interactions of transcription factors with the transcription machineries. These include for example, enhancer,^92^ silencer^93^ or switching elements,^94^ leading to temporal dynamic modulation of transcription and gene expression. For example, by coupling two mutually repressing transcription factor pathways, biomimetic oscillatory^95,96^ or bistable^97^ active gene expression were reported. Also, transient transcription machineries regulated by auxiliary enzymes^98^ or DNAzymes^99^ were reported. The allosteric ligand/aptamer transient activation of a transcription process is, to the best of our knowledge, unprecedented.

Figure 5A depicts schematically the transient Ade/ADA allosteric operation of a transcription machinery. The reaction module consists of the strands N_a_/T_a_ that exist as an inactive transcription template lacking a full T7 promoter sequence. The strand P_a_ contains the sequence x’ that hybridizes to the sequence x in strand N_a_ and completes the T7 promoter, however despite the complementarity of the domains x/x’ they are pre-engineered to form a non-stable five-base duplex. To assist the binding of P_a_ to the transcription template, Ade aptamer subunits, d_1_ and d_2_, were conjugated to strands P_a_ and N_a_. Added Ade cooperatively stabilize the formation of the active transcription template by the cooperative formation of the Ade/aptamer subunits complex and the x/x’ duplex completing the promoter domain in the transcription template. Formation of the intact Ade-stabilized transcription template activates, then, the T7 RNAP/NTPs transcription machinery transcribing the RNA product, R_2_. Since the affinity of the Ade aptamer towards other adenine-containing ligands is well established,^100^ ATP was excluded from the reaction circuit. The transcription template is pre-engineered to displace, by the transcribed RNA, R_2_, the auxiliary transducing F/Q DNA duplex that is comprised of a fluorophore-modified (FAM) strand and a quencher-modified (BHQ1) strand, where the fluorescent signal is effectively quenched, generating the fluorescent F/R_2_ duplex and the free quencher-modified strand Q. Thus, the time-dependent fluorescence changes in the system reflect the temporal performance of the transcription process induced by Ade. The ADA integrated in the circuit, temporally transformations Ade to inosine resulting in the depletion of Ade, leading to the separation of the strand P_a_ from the transcription template, that transiently recovers the parent reaction circuit. Thus, the ADA present in the system induces the transient dissipative evolution of the fluorescent intermediate F/R_2_ duplex in the system. The temporal fluorescence changes caused by the displacement induced by the RNA product reflect, then the dynamically modulated transcription occurring in the system. Figure 5B, panels I-III, depict the time-dependent fluorescence changes generated by the reaction module in the presence of different concentrations of Ade, yet in the absence of ADA: (i) 0 mM (ii) 0.125 mM (iii) 0.5 mM (iv) 2.0 mM. While in the absence of Ade minimal transcription of R_2_ occurs (Reflected by the lack of separation of the F/Q-transducing duplex), curve i, the time-dependent fluorescence changes are intensified as the concentration of Ade increases, curves ii-iv. Using an appropriate calibration curve relating the fluorescence of the released F-labeled DNA strand as a function of R_2_ concentration, Figure S6, the temporal transcribed RNA product R_2_ in the presence of different Ade concentrations is displayed in Figure 5B, panel II. Temporal first order maximum catalytic rates (V_max_) of the transcription template derived from panel II are displayed in panel III. The results confirm that Ade stabilizes allosterically the assembly of the promoter/transcription template P_a_/N_a_/T_a_ transcribing the product R_2_. Figure 5C depicts the temporal fluorescence changes generated by the transcription machinery, in the presence of variable Ade concentrations and a constant concentration of ADA = 0.025 U·ml^-1^. The temporal fluorescence changes, reflecting the rate of R_2_ production reveal a non-linear behavior tending to reach a saturation value. The temporal fluorescence changes and the resulting saturation levels are controlled by the concentration of Ade. As the concentration of Ade increases, the intensities of the fluorescence changes are higher, consistent with the increased concentrations of the Ade/aptamer subunits stabilized active transcription template. The non-linear temporal fluorescence intensity changes support that an accompanying mechanism slowing down the transcription process exists in the system, consistent with the ADA-induced depletion of the transcription machinery. Using a calibration curve relating the fluorescence intensities to the R_2_ RNA concentrations, Figure S6, the temporal concentrations of the transcribed R_2_ were evaluated, Figure 5C, panel II. The temporal catalytic rates (first order derivatives) of the transcription machinery, in the presence of different Ade concentrations, are displayed in panel III. Dissipative, transient catalytic rates revealing peak rates (V_max_) controlled by concentrations of Ade are observed (inset, panel III). Figure 5D, panel I depicts the temporal fluorescence changes associated with R_2_ transcription, in the presence of variable ADA concentrations, and a fixed concentration of Ade = 2.0 mM. As the concentration of ADA increases, the maximal fluorescence level is lower. Figure 5D, panel II shows the temporal concentration changes of the transcribed R_2_, and panel III displays the temporal catalytic rates of the transcription machinery. As the concentration of ADA increases V_max_ lowers and the dissipative depletion of the transcription process is faster (inset, panel III). These results are consistent with the faster depletion of the allosteric Ade/aptamer subunits stabilized transcription template, as the concentration of ADA increases.

**Figure 5.**
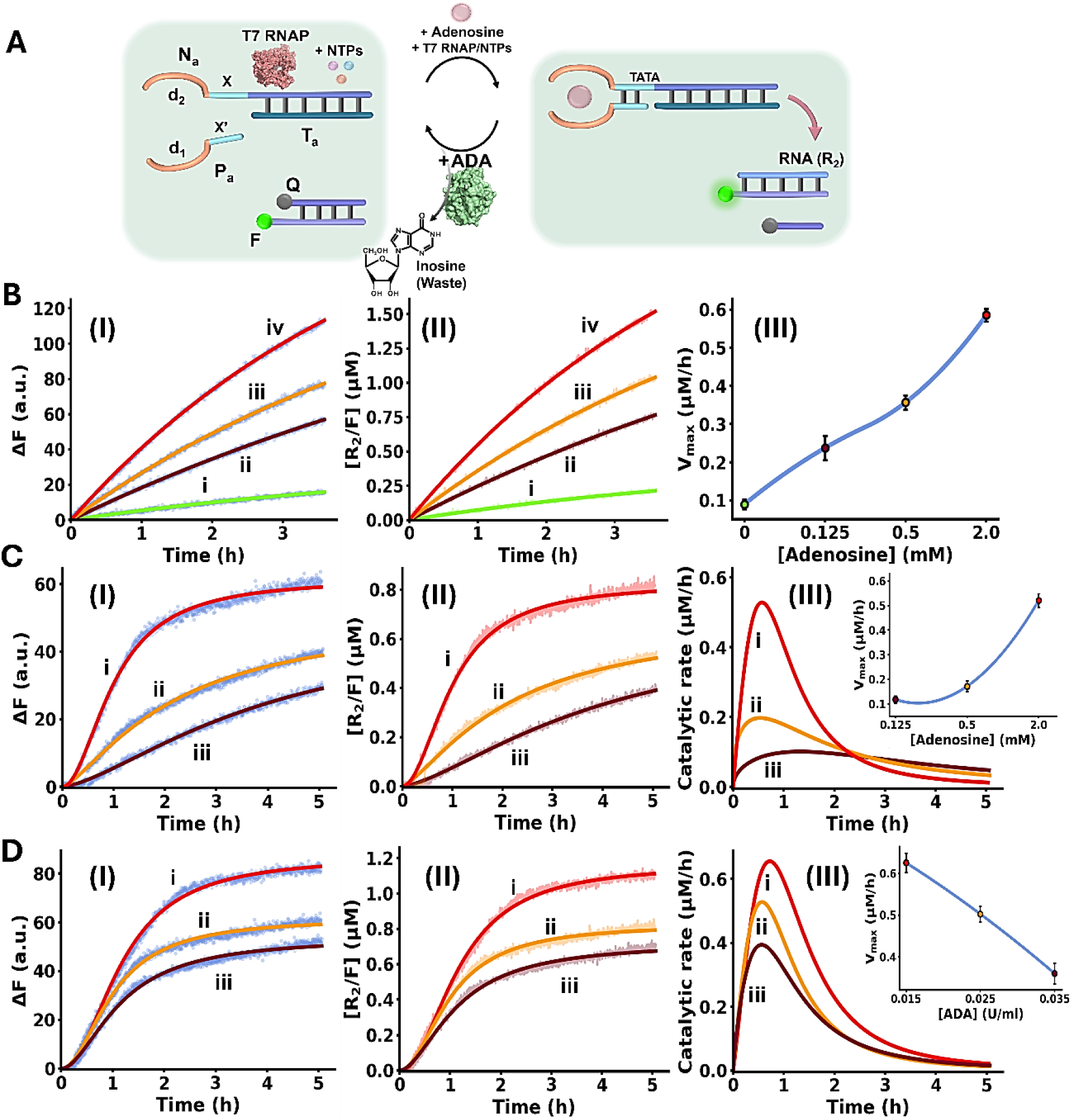
(A) Panel I – Schematic reaction module applying the Adenosine (Ade)/Adenosine deaminase (ADA) system for the transient allosteric operation of a transcription machinery emulating temporal transcription factor regulated transcription. (B) Temporal fluorescence changes generation by the Ade-driven operation of the reaction module, in the absence of ADA, and in the presence of variable Ade concentrations: (i) 0 mM, (ii) 0.125 mM, (iii) 0.5 mM, (iv) 2.0 mM. Panel II – Temporal concentration changes of R_2_, generated by the reaction module shown in (A), in the absence of ADA and in the presence of variable Ade concentrations: (i) 0 mM, (ii) 0.125 mM, (iii) 0.5 mM, (iv) 2.0 mM (Translation of the fluorescence changes shown in panel I into R_2_ concentrations, using the appropriate calibration curve, Figure S6). Panel III – Temporal catalytic rates corresponding to the generation of R_2_ by the reaction module, under variable Ade concentrations. (C) Panel I – Temporal fluorescence changes generated by the reaction module in the presence of ADA = 0.025 U·ml^-1^ and variable concentrations of Ade: (i) 2.0 mM, (ii) 0.5 mM, (iii) 0.125 mM. Panel II – Temporal concentration changes of R_2_ generated by the reaction module in the presence of ADA = 0.025 U·ml^-1^ and variable Ade concentrations: (i) 2.0 mM, (ii) 0.5 mM, (iii) 0.125 mM (Translation of the fluorescence changes shown in panel I into R_2_ concentrations, using the appropriate calibration curve, Figure S6). Panel III – Catalytic rates corresponding to the production of R_2_ in the presence of ADA = 0.025 U·ml^-1^, and variable Ade concentrations: (i) 2.0 mM, (ii) 0.5 mM, (iii) 0.125 mM (First order time dependent derivatives of the curves shown in panel II). Inset: Peak rates of transient formation of R_2_ at different Ade concentrations. (D) Panel I – Temporal fluorescence changes generated by the reaction module in the presence of Ade = 2.0 mM and variable concentration of ADA: (i) 0.015 U·ml^-1^, (ii) 0.025 U·ml^-1^, (iii) 0.035 U·ml^-1^. Panel II – Temporal R_2_ concentration changes in the presence of Ade = 2.0 mM and variable concentration of ADA: (i) 0.015 U·ml^-1^, (ii) 0.025 U·ml^-1^, (iii) 0.035 U·ml^-1^ (Conversion of the fluorescence changes displayed in panel I by the appropriate calibration curve, Figure S6). Panel III – Catalytic rates corresponding to the production of R_2_ in the presence of Ade = 2.0 mM, and variable ADA concentrations: (i) 0.015 U·ml^-1^, (ii) 0.025 U·ml^-1^, (iii) 0.035 U·ml^-1^ (First order time dependent derivatives of the curves shown in panel II). Inset: Peak rates of transient formation of R_2_ at different Ade concentrations. In all experiments, N_m_/T_m_ = 0.2 µM, P_m_ = 0.2 µM, NTPs = 3 mM, T7 RNAP = 1.2 U·µl^-1^. Data are means±SD, N=3.

## Conclusions

The study introduced allosteric ligand/aptamer complexes as functional structures orchestrating dynamic or transient catalytic DNA-based frameworks. These included the melamine (Mel)/aptamer subunits allosteric activation of the Mg^2+^-ion dependent DNAzyme, the allosteric inhibition of thrombin and the allosteric operation of a transcription machinery. Moreover, by coupling of the allosteric ligand/aptamer complex induced stabilization of the catalytic transformation to an auxiliary enzyme depleting the ligand, the transient, dissipative, operation of catalytic transformations was demonstrated. This was exemplified with the allosteric adenosine (Ade)/aptamer complex control over fibrinogenesis or transcription machineries and the transformation of these processes into dissipative and transient pathways in the presence of adenosine deaminase (ADA). Beyond mimicking native processes by synthetic circuits, such as transcription factor guided transcription machineries, the systems might be used for amplified sensing and biomedical applications, such as dose-controlled fibrinogenesis (blood coagulation) or a biomarker-induced synthesis of RNA inhibiting aptamers. The significance of the study is reflected by the versatility of the concepts. Many other ligand/aptamer subunits complexes may be envisaged to allosterically operate catalytic transformations, such as biomarker dictated cleavage of mRNAs. Also, the ligand/enzyme degradation pathways, such as acetylcholine^101^/acetylcholinesterase, uric acid^102^/uricase and xanthine^103^/xanthine oxidase may be applied to design other transient allosterically operating machineries. Moreover, at present, all systems were operated in homogeneous solutions. Integration of the circuits within liposomes could yield functional circuit-loaded synthetic cells-protocells,^104-106^ and fusion between the liposomes and native cells could provide versatile means to deliver the loads into the cells thereby signaling cell functions by artificial circuits.^91^

## Experimental Section

### Melamine-mediated stabilization of Mg^2+^-dependent DNAzyme subunits

DNAzyme reaction mixtures consisted of 1 µM of each DNAzyme subunits strands D_m_ and E_m_, 20 mM Tris acetate buffer pH 7.9, 40 mM MgCl_2_ and 2.25 µM BSA with varying concentrations of melamine. The mixture was incubated at room temperature for 1 hour. After the addition of 1 µM of the substrate strand, S, time-dependent fluorescence changes generated by the cleavage of the modified fluorophore/quencher substrate were monitored at 25°C (600V). The temporal concentrations of the free fluorophore were quantified using the calibration curve shown in Figure S1.

### Allosteric melamine-mediated inhibition of thrombin-induced fibrinogenesis

Reaction mixtures consisted of 1 µM of each of the DNA subunits in 40 mM Tris acetate buffer pH 7.9 and 40 mM MgCl_2_ with varying concentrations of melamine. Subsequent to a 1-hour incubation at room temperature, 5 nM thrombin was added to the solution and incubated at room temperature for 15 minutes. To evaluate thrombin coagulation activity, 10 mg/ml fibrinogen was added and the intensity changes of scattered light caused by formation of the fibrin network were immediately monitored at 25°C (500V)^[1]^.

### Transcription of the Malachite Green RNA aptamer from the allosteric melamine-triggered transcription machinery

For the melamine transcription experiments 20 µM of the template strands N_m_ and T_m_ were annealed in 1× RNAPol reaction buffer at 85°C for 5 minutes then cooled down to 25°C over 15 minutes.

A reaction mixture with varying concentrations of melamine, 0.2 µM annealed N_m_/T_m_, 0.2 µM P_m_ strand, 1× RNAPol reaction buffer, 20 mM MgCl_2_, 5 mM DTT, 0.5 mM NTPs and 2 µM MG was incubated at room temperature for 1 hour. The system was activated by the insertion of 1.5 U·µl^-1^ T7 RNAP, time-dependent fluorescence changes generated by the binding of MG/MG RNA aptamer were measured at 35°C (900V). The temporal concentrations of the malachite green RNA aptamer were quantified using the calibration curve shown in Figure S4.

### Allosteric adenosine/ADA-modulated, transient, dissipative, inhibition of thrombin-induced fibrinogenesis

The adenosine induced inhibition of fibrinogenesis reaction mixtures contained 1 µM of each of the DNA subunits in 40 mM Tris acetate buffer pH 7.9 and 40 mM MgCl_2_ was incubated for 1 hour at room temperature with variable concentrations of adenosine. Subsequent the incubation, 5 nM thrombin was added to the solution and incubated at room temperature for 15 minutes. To evaluate thrombin coagulation activity, 10 mg/ml fibrinogen was added and the intensity changes of scattered light caused by formation of the fibrin network were immediately monitored at 25°C (500V)^[1]^.

To probe the temporal activity of thrombin within the dissipative system containing ADA, a reaction mixture was prepared, one sample was withdrawn prior to the addition of adenosine. Next, adenosine was added and the reaction mixture was allowed to incubate for 1 hour at room temperature. Following incubation, variable concentrations of ADA were added, aliquots of 90µl were withdrawn from a reaction mixture at defined time intervals and incubated with 5 nM thrombin for 15 minutes at room temperature. To evaluate thrombin coagulation activity, 10 mg/ml fibrinogen was added and the intensity changes of scattered light caused by formation of the fibrin network were immediately monitored at 25°C (500V)[1].

### Transient, dissipative RNA transcription by the allosteric adenosine-activated, ADA-modulated, transcription machinery

For the adenosine-activated transcription experiments 20 µM of the template strands N_a_ and T_a_ were annealed in 1× RNAPol reaction buffer at 85°C for 5 minutes then cooled down to 25°C over 15 minutes. The fluorophore and quencher modified DNA strands were annealed following the same procedure.

A reaction mixture with varying concentrations of adenosine, 0.2 µM annealed N_a_/T_a_ strands, 0.2 µM P_a_ strand, 2 µM annealed F/Q strands, 1× RNAPol reaction buffer, 20 mM MgCl_2_, 5 mM DTT and 3 mM NTPs was incubated at room temperature for 1 hour. The system was activated by the insertion of 1.2

U·µl^-1^ T7 RNAP, time-dependent fluorescence changes generated by the displacement of the quencher strand by the RNA product were measured at 33°C (600V).

Dissipative transcription experiments with ADA were prepared and measured in the same manner. Following activation by T7 RNAP, varying concentrations of ADA were added to the system incubated with 2 mM adenosine or alternatively, 0.025 U·ml^-1^ ADA was added to the system incubated with different concentrations of adenosine. The temporal concentrations of the RNA generated by transcription machinery were quantified using the appropriate calibration curve shown in Figure S6.

### ITC experiments

ITC reaction solutions were prepared in 20 mM Tris acetate pH 7.9 and 40 mM MgCl_2_ (Total volume 300µl). Each experiment consisted of 17 injections of 2.3µl adenosine, melamine or analog molecules at 5 s time intervals into DNA aptamer solutions ranging from 10 to 20 µM, stirred at 750 rpm and held at 25°C. A delay of 150 s between injections was allowed for equilibration. All experiments were corrected for the heat of dilution of the titrant. Data was fit to a one set of binding sites binding model using MicroCal PEAQ-ITC Analysis Software. Measurement was performed using a MicroCal PEAQ-ITC instrument (Malvern).

## Supporting Information

Supporting Information Available: Chemicals, oligonucleotide sequences, calibration curve corresponding to the Mel-assisted DNAzyme activity, ITC experiments, table summarizing ITC information, thrombin activity in the presence of Mel or Ade, calibration curves corresponding to RNA transcription by allosteric transcription machineries, scheme illustrating dissipative processes.

## Present Addresses

**Jiantong Dong** - State Key Laboratory of Medicinal Chemical Biology, Tianjin Key Laboratory of Biosensing and Molecular Recognition, Research Center for Analytical Sciences, College of Chemistry, Nankai University, Tianjin 300071, P. R. China

## Author Contributions

All authors have given approval to the final version of the manuscript.

## Notes

The authors declare no competing financial interest.

## Abbreviations

Mel: Melamine
Ade: Adenosine
ADA: Adenosine deaminase
ITC: Isothermal titration calorimetry

