## Supplemental Information for "Allosteric Ligand-Aptamer Complexes Orchestrate Supramolecular or Transient Catalytic, Transcription and Fibrinogenesis Processes"

### Materials and Methods

#### Materials

DNA oligonucleotides, substrate/quencher-modified substrates and RNA were purchased through Integrated DNA Technologies Inc. T7 RNA polymerase (50,000 units/mL, 0.8 μM), ribonucleotide (NTP) Mix (GTP, ATP, UTP and CTP, each 25 mM), separated ribonucleotides (UTP, GTP, CTP, each 100mM) and 10 × RNAPol reaction buffer (400 mM Tris-HCl, 60 mM $MgCl_2$, 10mM DTT, 20 mM spermidine, pH 7.9 @ 25 °C) were purchased from New England BioLabs Inc. Thrombin from human plasma (≥ 2,000 NIH units/mg protein, MW = 37.4 kDa), fibrinogen from human plasma (50–70% protein), adenosine deaminase (1,000 units/mL), melamine, adenosine, $MgCl_2$, tris acetate, BSA, DTT, RNAse free water and Malachite Green (MG) were purchased from Sigma-Aldrich.

#### Instrumentation

DNA concentrations were determined by UV-1900 spectrophotometer (Shimadzu). Time dependent fluorescence changes were recorded with a Cary Eclipse Fluorometer (Agilent Technologies) using a quartz cuvette with 10-mm path length supplied by Hellma Analytics, FAM was excited at 496 nm and emission was recorded at 520 nm. Malachite green was excited at 632 nm and emission was recorded at 650 nm. Light scattering changes were recorded at 650 nm using a plastic cuvette with 10-mm path length purchased from Brand GMBH, Wehrheim, Germany. Isothermal titration calorimetry measurements were performed on the PEAQ-ITC instrument (Malvern).

**Table S1.** Nucleic acid strands used in this study

| Name | Sequence (5' – 3') |
|---|---|
| $E_m$ | 5′-ACCTTTAGGGGGTGTGC**CACCCATGT**TCCTGA-3′ |
| $D_m$ | 5′-CTGTTC**AGCGAT**GCACACCGATGGCGGTCCTGTAGGT-3′ |
| S | 5’-FAM-TCAGGATrAGGAACAG-BHQ1-3’ |
| $G_m$ | 5′-AGGGTGGTGGCGAAAGCACACCGATGGCGGTCCTGTAGGT-3′ |
| $H_m$ | 5′-ACCTTTAGGGGGTGTGCAAACGCCTAGGTTGGGT-3′ |
| $N_m$ | 5’-TAATACGACTCACTATAGGGATCCCGACTGGCGAGAGCCAGGTAAC GAATGGATCC-3’ |
| $T_m$ | 5’-GGATCCATTCGTTACCTGGCTCTCGCCAGTCGGGATCCCTATAGTG AGTCTTAGGGGGTGTGC-3' |
| $P_m$ | 5’-GCACACCGATGGCGGTCCTGTGTATTA-3' |
| $R_1$ | 5′- GGAUCCCGACUGGCGAGAGCCAGGUAACGAAUGGAUCC -3′ |
| $G_a$ | 5′-AGGGTGGTGGCGAACTGCGGAGGAAGGT-3′ |
| $H_a$ | 5′-ACCTGGGGGAGTATGATCGCCTAGGTTGGGT-3′ |
| $N_a$ | 5’-ACCTGGGGGAGTATTAATACGACTCACTATAGGGTTCTCCTCTGCG TTGTTGTTTG-3’ |
| $T_a$ | 5’-CAAACAACAACGCAGAGGAGAACCCTATAGTGAGTCG-3' |
| $P_a$ | 5’-TATTATGCGGAGGAAGGT-3’ |
| $R_2$ | 5′- GTTCTCCTCTGCGTTGTTGTTTG -3′ |
| F | 5′-FAM-CAAACAACAACGCAGAGGAGAAC-3′ |
| Q | 5′-GTTCTCCTCTGCGTT-BHQ1-3′ |
| Intact Mel-ITC | 5'-GCACACCGATGGCGGTCCTGTTTAGGGGGTGTGC-3' |
| Split Mel-ITC | (a) 5'-GCACACCGATGGCGGTCCTGTAGGT-3'<br>(b) 5'-ACCTTTAGGGGGTGTGC-3' |
| Note | DNAzyme subunits are marked in bold. The ribonucleotide modification is colored in red. The T7 promoter is colored in blue and thrombin aptamer subunits are colored in orange. |

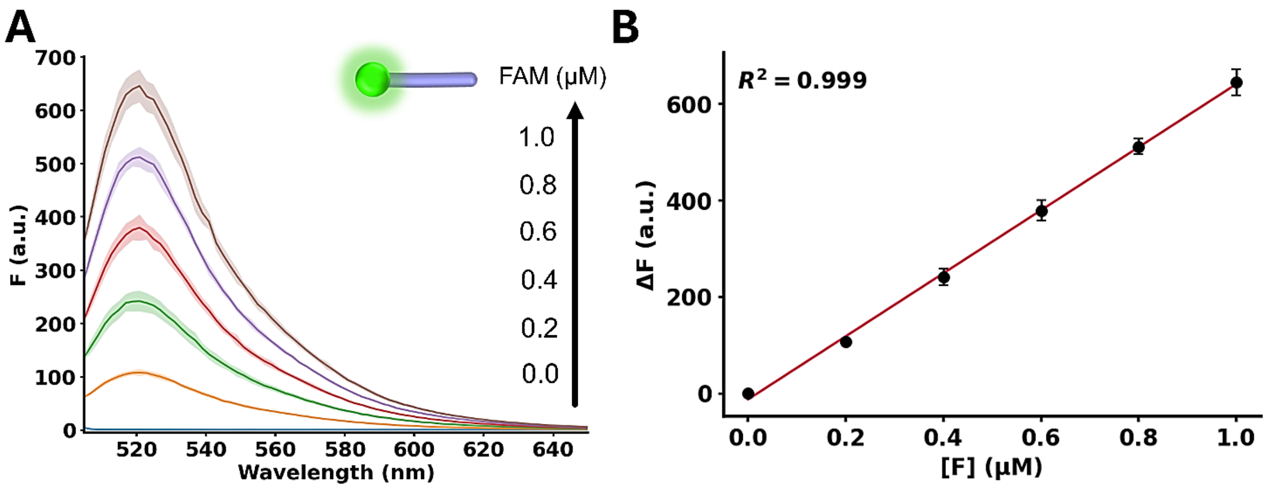


**Figure S1.** (A) Fluorescence spectra of different fluorophore (FAM) concentrations ($\lambda_{ex}$= 496 nm). (B) Derived calibration curve corresponding to the fluorescence intensities increasing as a function of the fluorophore concentration at $\lambda_{em}$= 520 nm. $R^2$=0.999.

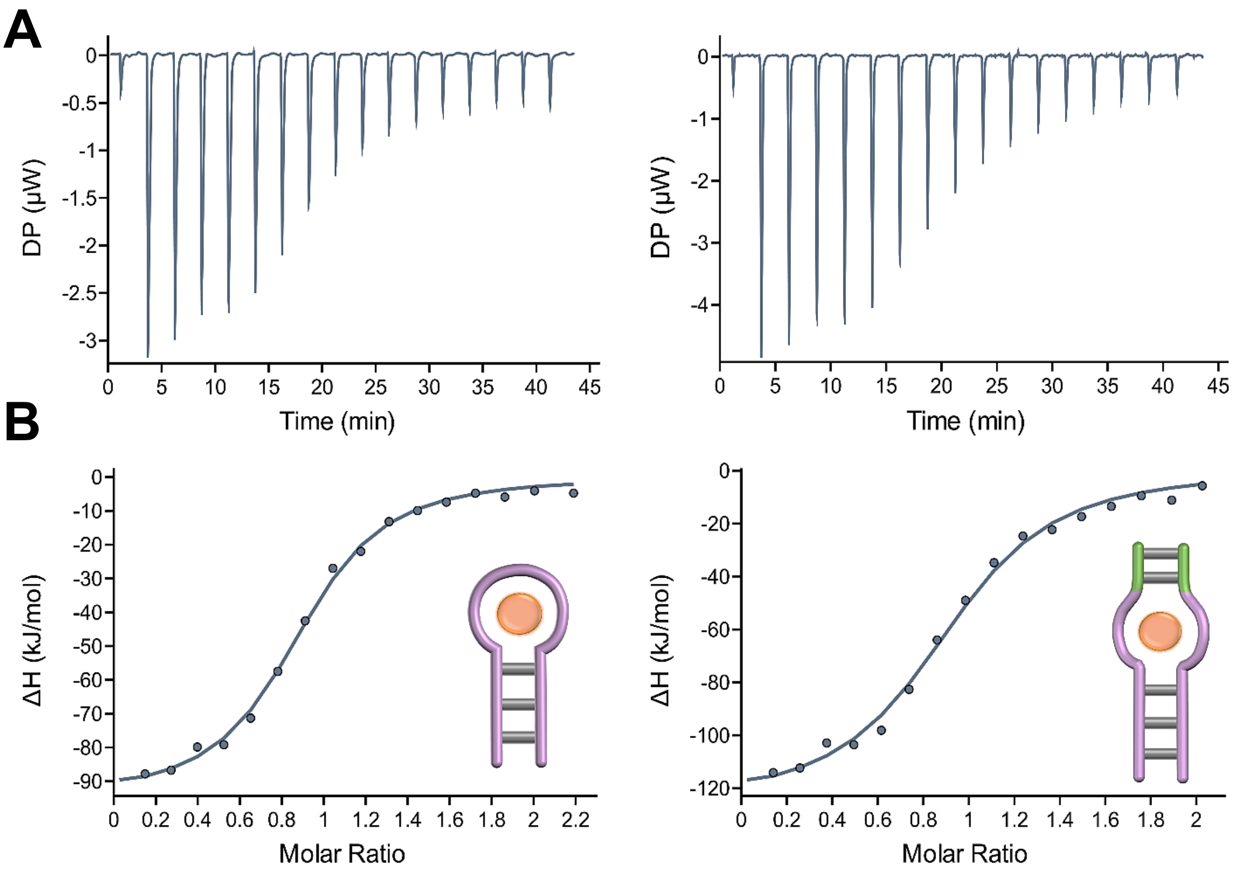


**Figure S2.** Isothermal titration calorimetry (ITC) experiments determining the $K_d$, stoichiometry and thermodynamics of the binding of melamine to the intact melamine aptamer and the split melamine aptamer. (A) Thermograms of melamine titrations into an intact melamine aptamer (left) and split melamine aptamer subunits (right). (B) Integrated curves representing the total heat exchanged derived from (A), intact aptamer (left) and split aptamer subunits (right).

**Table S2.** Isothermal titration calorimetry (ITC) analysis of Mel/aptamer

| **DNA** | **$K_d$** (μM) | **N** (sites) | **ΔG** (kJ·mol$^{-1}$) | **ΔH** (kJ·mol$^{-1}$) | **-TΔS** (kJ·mol$^{-1}$) |
|---|---|---|---|---|---|
| Intact aptamer | 0.800 ±0.139 | 0.869 ±0.02 | -34.8 | -95.1 ±2.89 | 60.3 |
| Split aptamer | 0.856 ±0.115 | 0.902 ±0.017 | -34.7 | -125 ±3.15 | 90.3 |

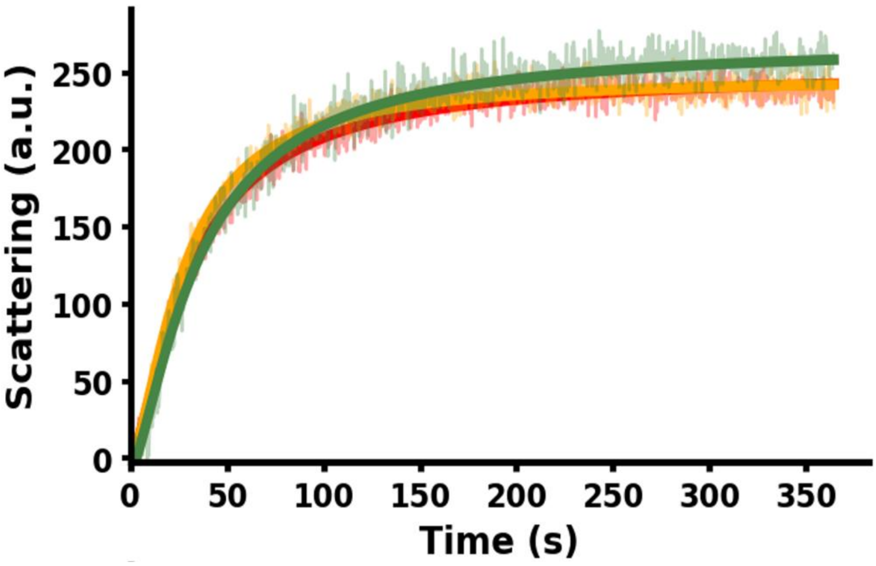


**Figure S3.** Temporal light scattering probing thrombin activity in the absence of DNA without additives and with 2 mM of adenosine or melamine. Samples were analyzed following the procedure described previously. Data are means±SD, N=3.

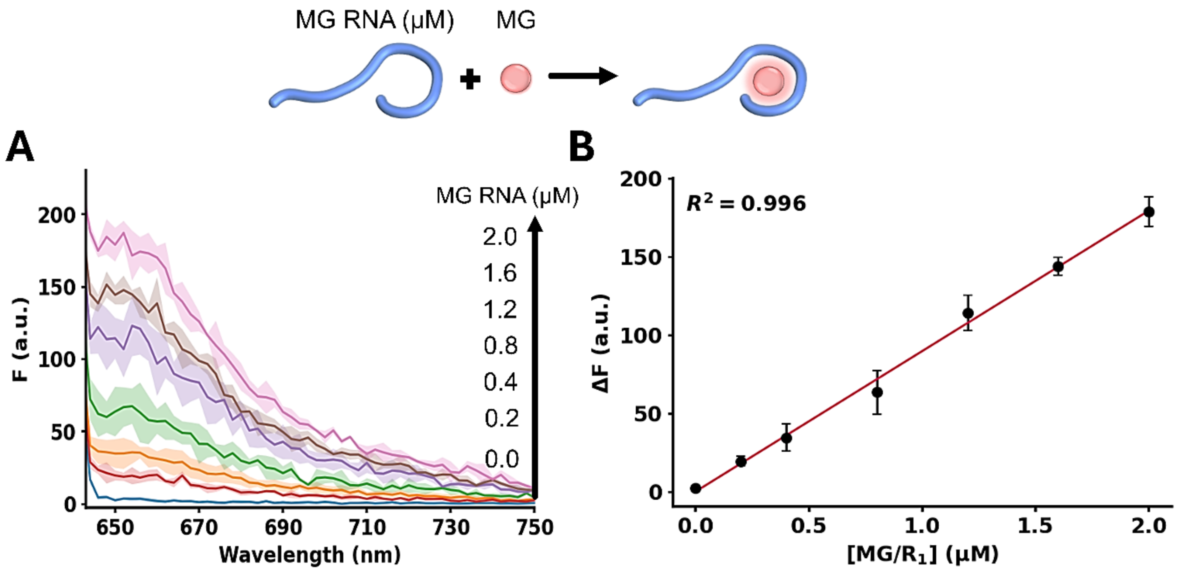


**Figure S4.** (A) Fluorescence spectra obtained from the Malachite Green (MG)/RNA aptamer complex, in the presence of variable concentrations of the RNA aptamer ($\lambda_{ex}$= 632 nm). (B) Derived calibration curve corresponding to the fluorescence intensities of the MG/RNA aptamer complex as the concentration of the malachite green RNA aptamer increases at $\lambda_{em}$= 650 nm. $R^2$=0.996.

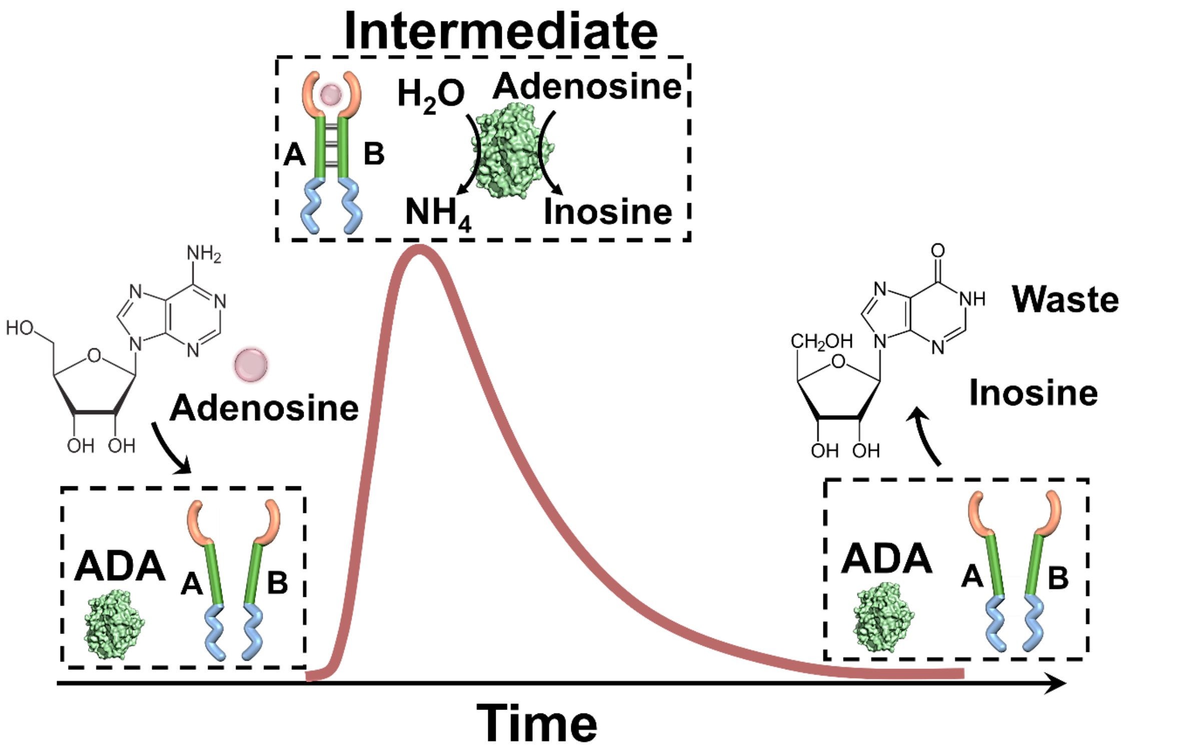


*** The concept is applicable to other dissipative ligand/aptamer complexes coupled to ligand degrading enzymes such as uric acid/uricase and acetylcholine/acetylcholinesterase.**

**Figure S5.** General scheme of the dissipative transient operation of a DNA-based reaction circuit including the split adenosine aptamer subunits present in strands A and B that are unable to form a stable duplex. Addition of adenosine activates the system, creating the intermediate product Ade/A/B interstrand complex. However, adenosine deaminase (ADA) present in the reaction module, deaminates adenosine to inosine, which lacks affinity to the adenosine aptamer subunits, leading to the separation of A and B and the dissipative deactivation of the system.

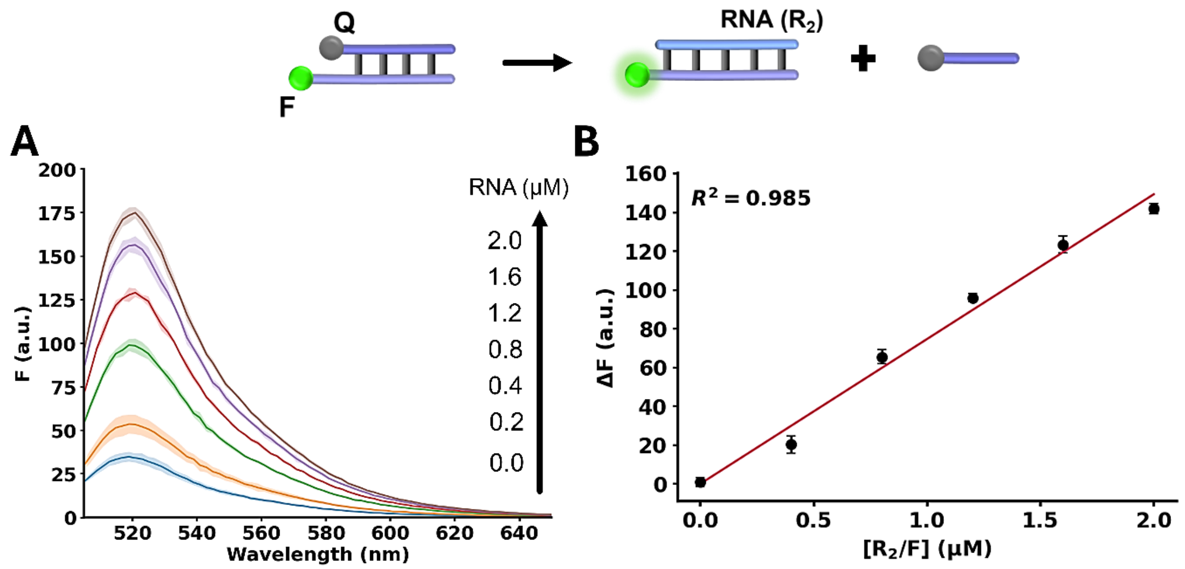


**Figure S6.** (A) Fluorescence spectra of FAM-labeled F DNA strand (2.0 µM) generated by the addition of different concentrations of $R_2$ resulting in the displacement of the BHQ1-modified Q DNA strand ($\lambda_{ex}$= 496 nm) from the F/Q duplex. (B) Derived calibration curve corresponding to the fluorescence intensities increasing as the concentration of $R_2$ increases at $\lambda_{em}$= 520 nm. $R^2$=0.985.
